# Association of Sonic Hedgehog with the extracellular matrix requires its zinc-coordination fold

**DOI:** 10.1101/2019.12.17.880039

**Authors:** Carina Jägers, Henk Roelink

## Abstract

Sonic Hedgehog (Shh) has a catalytic cleft characteristic for zinc metallopeptidases and has significant sequence similarities with some bacterial peptidoglycan metallopeptidases defining a subgroup within the M15A family that, besides having the characteristic zinc coordination domain, can bind two calcium ions. Extracellular matrix (ECM) components in animals include heparan-sulfate proteoglycans, which are analogs of bacterial peptidoglycan and thus potentially involved in the extracellular distribution of Shh. We found that the zinc-coordination fold of Shh is required for its association with ECM as well as for non-cell autonomous signaling. Association with the ECM requires the presence of at least 0.1 μM zinc and is prevented by mutations affecting critical conserved catalytical residues as well as extracellular calcium. Our results demonstrate that the zinc-coordination fold is required for ECM-association and suggest that the putative intrinsic peptidase activity of Shh is required for non-cell autonomous signaling.

## Introduction

The *Hedgehog* (*Hh*) gene was first identified in the *Drosophila melanogaster* screen performed by Christiane Nüsslein-Volhard and Eric Wieshaus in the late 1970s (Nüsslein-Volhard and Wieshaus, 1980). Like other segment polarity genes found in this screen, *Hh* genes are widely conserved among animals, and mammals have three Hh paralogs (Sonic -, Indian -, and Desert Hedgehog) that play roles in development (Echelard et al., 1993). Like all other Hhs, Shh is synthesized as a pro-protein that undergoes autoproteolytic cleavage mediated by the C-terminal part yielding an N-terminal part (ShhNp) that is the active ligand. Structural analysis of ShhN revealed its similarity to zinc-peptidases and Shh coordinates a zinc ion with residues H141, D148, and H183 (Hall et al., 1995). The notion that Shh signaled through a peptidase activity was quickly rejected by demonstrating that mutation of a critical residue involved in catalysis (E177) did not impair the ability of Shh to activate the Hh response (Fuse et al., 1999), and consequently the zinc coordination domain of Shh is often referred to as its “pseudo active” site (Bosanac et al., 2009; Maun et al., 2010). Still, a role for the zinc coordination fold is supported by the finding that Shh-E177A is unable to mediate non-cell autonomous long-range signaling from the notochord to the overlying neural plate (Himmelstein et al., 2017). Perhaps unsurprisingly, the zinc-coordination domain is found mutated in some individuals with the Shh signaling-related birth defect holoprosencephaly (Roessler et al., 1996; Traiffort et al., 2004), further indicating that the zinc-coordination fold of Shh is important for normal function. This is consistent with structures of Shh complexed with its receptor Patched1 (Ptch1), showing that the N-terminal 22 residues of Shh that are not part of the zinc-coordination fold, mediate binding to Ptch1 (Gong et al., 2018; Qi et al., 2018a; Qi et al., 2018b) and suffice to regulate Ptch1 activity (Tukachinsky et al., 2016).

Some bacterial species have conserved genes coding for peptidases that coordinate zinc and calcium identically to Shh (Rebollido-Rios et al., 2014; Roelink, 2018). These bacterial peptidases (members of the M15A subfamily of zinc D-Ala-D-Ala carboxypeptidases) cleave murein peptidoglycans, suggesting that Shh too might cleave a glycan-modified protein, possibly a matrix heparan sulfate proteoglycan (HSPGs). HSPG are an integral part of the extracellular matrix (ECM) and play in important role in the transport and presentation of several morphogens, including Hhs (Yan and Lin, 2009). Several HSPGs bind Shh and can both negatively and positively affect the Shh response (Capurro et al., 2008; Carrasco et al., 2005; Guo and Roelink, 2019; Witt et al., 2013). Furthermore, mutations in *Ext1* and *-2* coding for glycosyltransferases that catalyze glycosaminoglycan addition to the core proteins, disrupt Hh signaling in vertebrates (Guo and Roelink, 2019; Siekmann and Brand, 2005) and insects (Bellaiche et al., 1998).

By mutating residues in the zinc-coordination fold that are conserved between bacterial Hh-like peptidases and Shh, we provide evidence that this domain is required for the association of ShhN to the ECM and for non-cell autonomous signaling. Release of Shh into the ECM is enhanced in the presence of μM amounts of zinc indicating that this ion is an agonist of Shh. The ECM-associated Shh is active in signaling, indicating that the zinc-coordination fold of Shh mediates its release into the ECM to facilitate non-cell autonomous signaling, possibly through an intrinsic metallopeptidase activity of Shh.

## Results

### ShhN associates with the extracellular matrix

Due to a high sequence similarity to bacterial murein peptidases (Roelink, 2018), a conceivable function for the Shh zinc-coordination fold could be to modify proteoglycans thus affecting its extracellular matrix (ECM) association. We assessed ECM-bound Shh in the fraction of macromolecules that remain on the tissue culture plate after non-lysing cell removal. Hek293t cells were transfected with *Shh* (mutant) constructs and after two days removed by washing with PBS and mild agitation. In order to exclude a role for Dispatched1 in the association of Shh with the ECM (Tian et al., 2005), we used Shh-C199* (ShhN) and found that ShhN could readily be detected in the ECM fraction using gel electrophoresis followed by SYPRO Ruby staining (Figure 1A). Visualizing Shh-C199* by staining of the decellularized plates with the anti-Shh mAb5E1, well-defined Shh “footprints” of Shh producing cells were observed (Figure 1B). Shh-C199* is commonly referred to as a “soluble” protein due to the absence of C-terminal cholesterol and its well-established presence in the supernatant of Shh-producing cells. The distinct Shh “footprints”, however, indicate that the protein leaves the cell and enters the adjacent ECM in a direct manner and not via an intermediate soluble form from the medium which would result in a more homogeneous Shh distribution in the ECM. Surprisingly, more Shh could be detected in the ECM than in the lysate of *Shh-C199\** transfected Hek293t cells (Figure 1B), together showing that entry of ShhN into the ECM is robust. Shh-responsive LightII cells plated on the decellularized and Shh-conditioned ECM showed that it is able to elicit a transcriptional Hh pathway response similar to that of ShhN-conditioned medium (Figure 1C).

**Figure 1:**
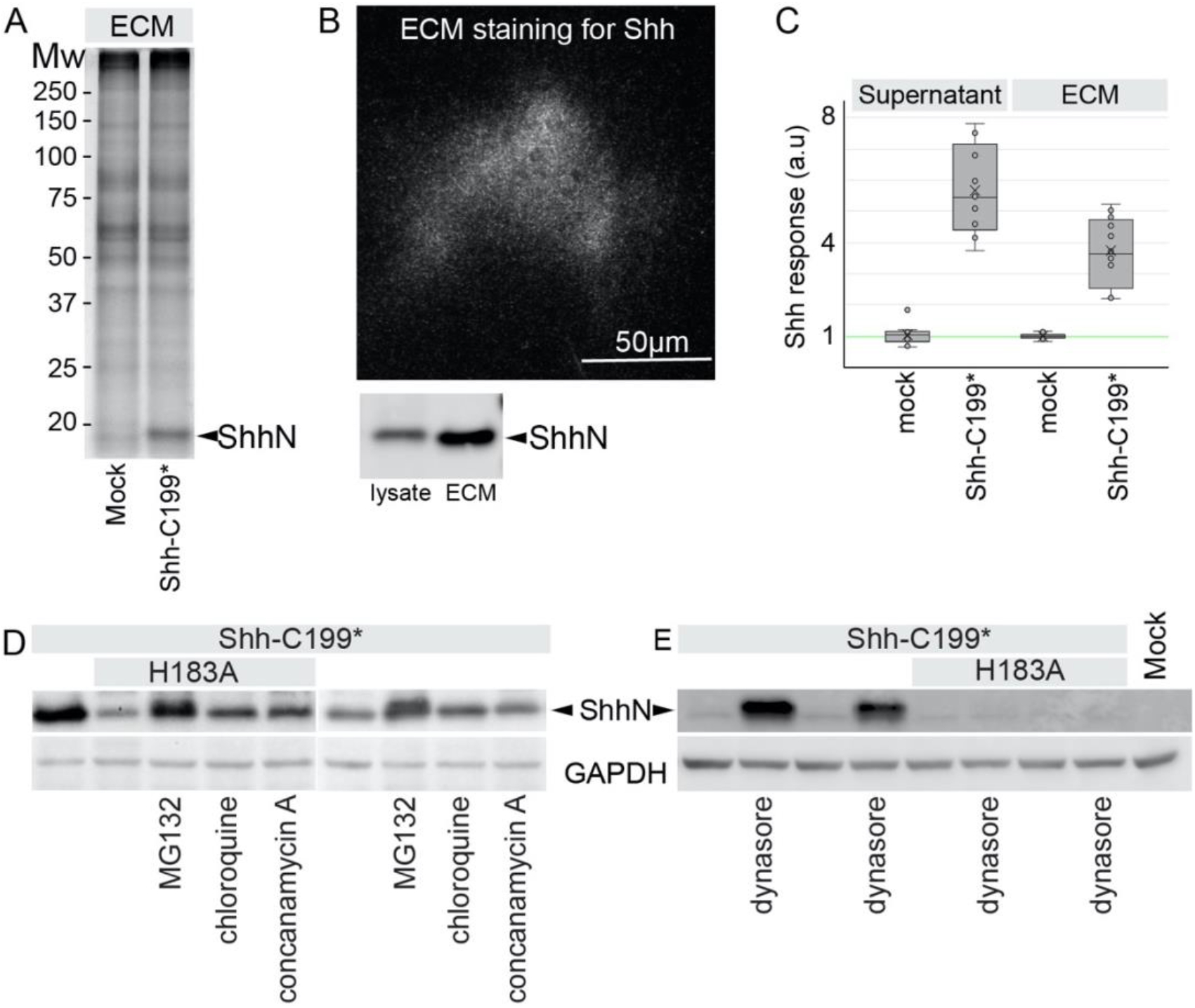
Active ShhN associates with the extracellular matrix. **A:** ECM deposited by mock and *Shh-C199\**-transfected Hek293t cells analyzed by SDS-PAGE and SYPRO-Ruby staining. ShhN is indicated. **B:** *Shh-C199\**-transfected Hek293t cells were plated on glass slides and removed after 24h. The slides were stained with mAb5E1, showing the presence of ShhN. Scale bar is 50μm. **C:** supernatant and ECM conditioned by *Shh-C199\**-transfected Hek293t cells. LightII cells were either grown on mock or ShhN conditioned ECM. Cells grown on ECM deposited by mock transfected cell were grown in the absence or presence of mock or Shh-C199* conditioned supernatant. Box and whisker plots, n ≤ 3. **D:** Western blot analysis of HEK293t cells transfected with the indicated Shh mutants. 100 nM MG-132 (proteasome inhibitor), 100 nM Chloroquine and 100 nM Concanamycin A (inhibitors of endosome acidification) were assessed for their ability to affect Shh accumulation. **E:** Western blot analysis of HEK293t cells transfected with the indicated Shh mutants, and the effects of the dynamin inhibitor Dynasore (50 μM) was assessed for its effect on Shh accumulation.

### Mutations in the Zinc-coordination domain reduce the stability of Shh-C199*

Mutating the residues directly involved in the coordination of zinc are obvious candidates to assess a role for the zinc-coordination fold. However, Shh mutants in the zinc coordination domain could barely be detected as the N-terminal processed form (ShhNp) on Western Blots, despite normal detection of the Shh pro-protein (Casillas and Roelink, 2018). We and others (Traiffort et al., 2004) initially incorrectly interpreted this as a failure of auto-processing, but the same low levels were observed analyzing these mutants in Shh-C199*, a Shh mutant truncated at the auto-processing site (ShhN) (Roelink et al., 1995). Addition of the proteasome inhibitor MG132 (1996) or the inhibitor of endosome acidification by chloroquine resulted in ShhN accumulation of Shh-C199*/H183A (ShhN/H183A), possibly indicating a misfolded protein-induced degradation of this mutant (Figure 1D). The Dynamin inhibitor Dynasore (that inhibits endocytosis) (Macia et al., 2006) causes strong accumulation of Shh-C199*, but not of Shh-C199*/H183A, further indicating that the destabilization of Shh-C199*/H183A occurs before it reaches the plasma membrane (Figure 1E). We found that other zinc coordination mutations as well as several holoprosencephaly-associated point mutations in Shh cause its destabilization, indicating a role for increased ShhN degradation in this birth defect (Casillas and Roelink, 2018). In general, we will not use these mutants with a reduced half-life.

### The zinc-coordination fold of Shh is required for association with the ECM

We hypothesized that metal occupancy of the zinc-coordination domain is required for normal Shh function. The Kd for zinc binding to Shh in the absence of calcium appears to be low (Day et al., 1999), but DMEM tissue culture medium has no added zinc and is thus expected to have only small amounts of this ion. While the amount of protein in the lysate of Hek293t cells transfected with *Shh-C199\** remained relatively unchanged with increasing zinc concentrations, the amount of ECM-bound Shh-C199* increased several-fold with an EC_50_ between 0.1 and 1 μM zinc (Figure 2 B-F). This indicates that there is little effect of zinc on Shh synthesis and intracellular stability, but that occupancy of the zinc coordination fold enhances ECM association. This effect on ShhN ECM accumulation was specific to zinc as other divalent cations like copper and magnesium failed to increase the amount of ShhN in the ECM (Figure S2).

**Figure 2:**
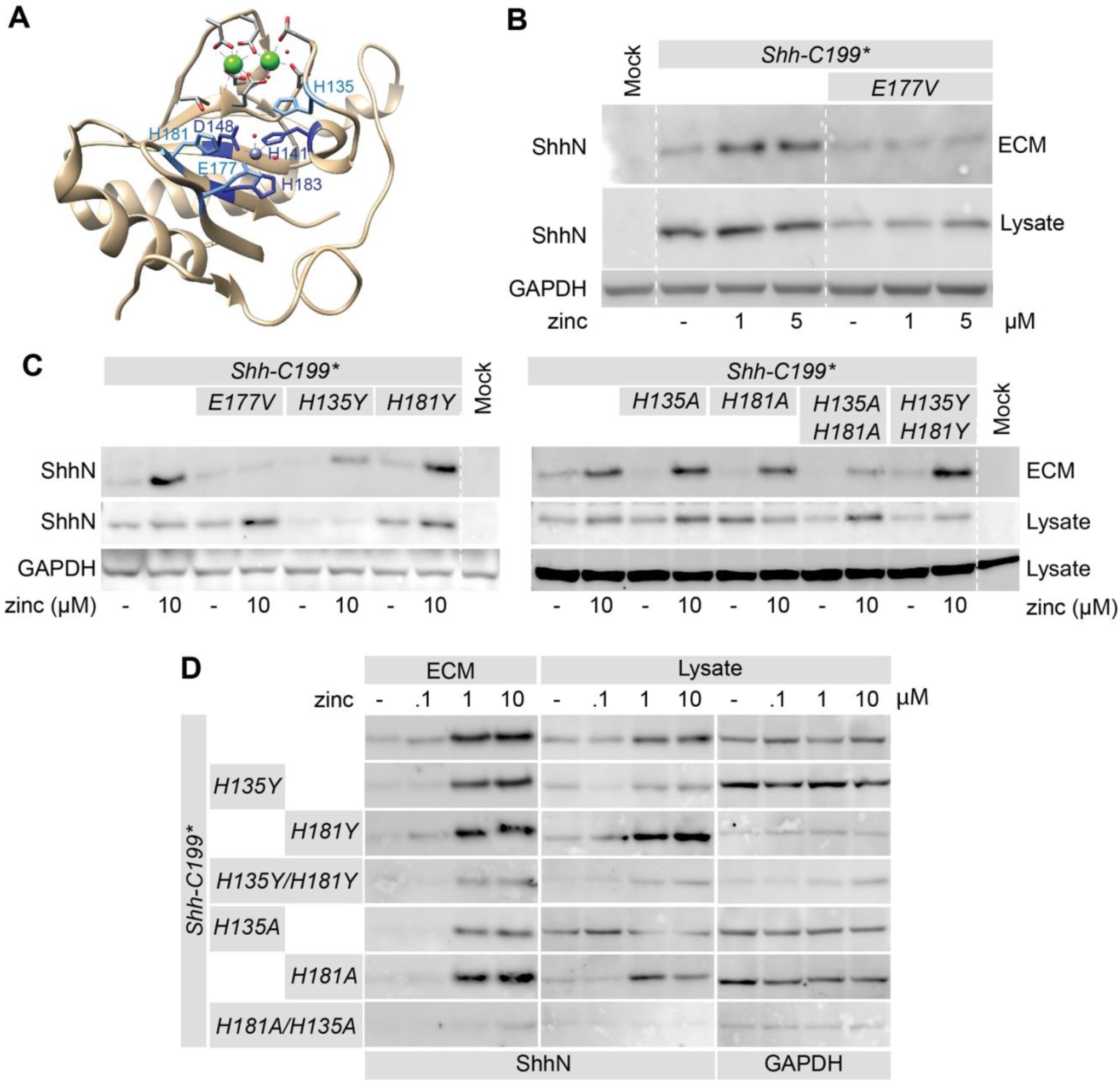
ECM-association of ShhN requires zinc in and putative peptidase activity. **A:** Diagram of the Shh structure (3D1M) with relevant residues indicated. **B:** Western blot analysis of the lysate and ECM of HEK293t cells transfected with *Shh-C199\** and *Shh-C199*/E177V* and cultured in DMEM containing 0.18 mM calcium and the indicated concentrations of zinc. **C:** The effect of mutations of the transition state-stabilizing residues H135 and H181 to alanine (A) or tyrosine (Y) on the zinc-dependent accumulation in the ECM was analyzed on a Western Blot of the extracted ECM from transfected HEK293t cells cultured in 0.18 mM calcium with or without 10 μM zinc. **D:** zinc dose-response analysis of H135 and H181 mutations assessed by Western blot of the lysate and ECM of HEK293t cells transfected with the indicated mutants and cultured in 0.18 mM calcium and increasing concentrations of zinc (0.1, 1, 10 μM).

Besides the zinc coordinating residues, the glutamic acid residue at position 177 (E177, mouse numbering) is well-conserved and in close proximity to the zinc-coordinating residues. In Shh-related peptidases the E177 equivalent is required for catalytic activity. The mutants Shh-C199*/E177A and -/E177V are predicted to be able to coordinate zinc but could reveal a function of the zinc-coordinating fold. Unlike the zinc coordination mutants, we found the Shh-C199*/E177A and -/E177V mutants to be stable in the lysate (Figure 2B). The amount of Shh-C199*/E177V in the ECM was similar to that of Shh-C199* under low zinc concentrations but failed to further accumulate in the ECM under increasing zinc concentrations, demonstrating that ShhE177 is required for ECM association (Figure 2B), and that the zinc effects are not primarily mediated by an Shh-independent zinc sensitive event These observation supports the notion that a catalytic activity intrinsic to Shh is required for its association with the ECM.

### Mutating residues in the zinc-coordination fold of Shh affect association with the ECM

A second group of conserved residues in the zinc-coordination fold are two histidine residues with stacking sidechains, H135 and H181. These two histidine residues are conserved between Shh and BacHhs, but either one can be a tyrosine residue in M15A peptidases, and a tyrosine residue is present in the position homologous to H181 in butterfly and moth Hhs (e.g. NCBI PCG69308.1). We mutated either or both histidine residues 135/181 into alanine or tyrosine residues (*Shh-C199*/H135Y*or*A*, *Shh-C199*/H181Y*or*A*) and found that these forms of Shh process normally and are stable in the lysate (Figure 2C). Mutants with one or two tyrosine substitutions as well as single alanine substitutions were found at lower levels in the ECM but could be rescued under high zinc conditions (1, 10 μM), suggesting that tyrosine residues are to some extent synonymous mutations. Only *Shh-C199*/H135A/H181A* poorly associated with the ECM in the presence of zinc. As H135 and H181 could affect zinc binding of Shh, we tested if these mutants had an altered EC_50_ for zinc. We found that all mutants have a similar EC_50_ (Figure 2D), consistent with the notion that these residues are not directly involved in zinc coordination. Together with *Shh-C199*/E177V*, our findings using *Shh-C199*/H135A/H181A* further support the notion that the zinc-coordination fold of Shh is required for ECM association.

### Shh-C199* mutants unable to bind calcium remain sensitive to zinc

The overall structure of ShhN and the BacHhs indicate that they consist of a regulatory calcium-binding and a catalytic zinc coordinating domain (Rebollido-Rios et al., 2014), making up most of ShhN outside the extreme N-terminal Ptch1-binding domain. With the exception of BacHhs, bacterial M15A metallopeptidases lack the Hh/BacHh-type calcium coordination domain, and this domain is thus unlikely to be required for catalytic function *per se*. We made a Shh-C199* mutant that lacked all calcium-coordinating residues (Shh-C199*/E90A/E91D/D96A/E127A/D130N/D132L, Shh-C199*-Ca_Free_) and this form of Shh should be unable to bind calcium. After transfection, more ShhN was detected in lysates of cells cultured in the presence of higher calcium levels, but that was also observed in the Shh-C199*-Ca_Free_ expressing cells, and thus unlikely a direct effect of calcium on Shh. Increased amounts of ShhN in the lysate at higher calcium concentrations complicated the interpretation of the effects of calcium on ShhN accumulation in the ECM. However, while ShhN accumulation in the ECM varied with calcium concentrations, that of the Shh-C199*-Ca_Free_ mutant remained at the same level, indicating that this mutant is insensitive to extracellular calcium as measured by ECM association (Figure 3A).

**Figure 3:**
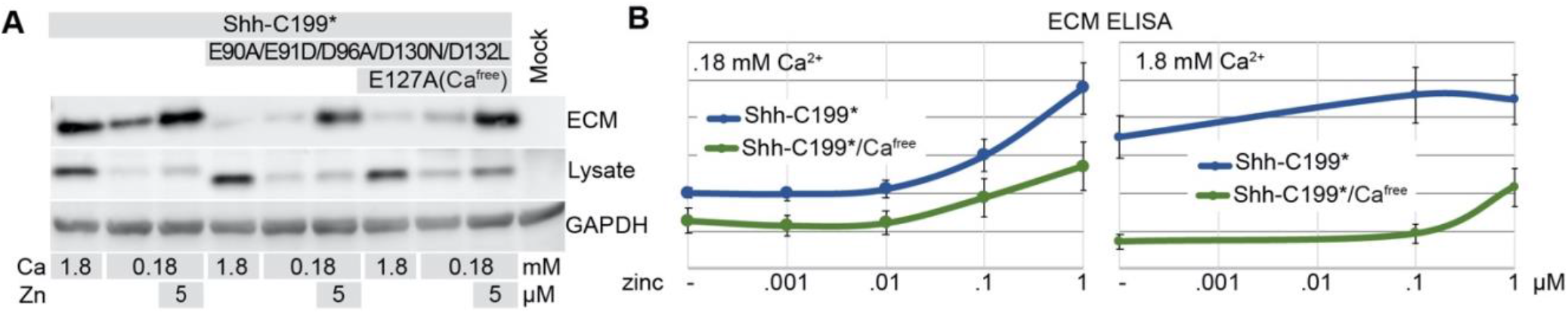
Calcium alters the sensitivity of Shh to zinc. **A:** Western blot analysis of lysates and ECM of HEK293t cells transfected with *Shh-C199\** and *Shh-C199*/E90A/E91D/D96AD130N/D132L*, or *Shh-C199\** and *Shh-C199*/E90A/E91D/D96A/E127A/D130N/D132L* (*Ca_Free_*), cultured in the presence of 0.18 or 1.8 mM calcium, and in the absence or presence of 5 μM added zinc. **B:** ECM-associated Shh-C199*N or Shh-C199*/Ca_Free_ deposited by transfected HEK293t cells was assessed by ELISA in the presence of .18 mM calcium (left panel) or 1.8 mM calcium (right panel) and increasing zinc concentrations as indicated. Shown are means and standard errors, n=6.

One possible mechanism of calcium regulating the transition from a cell-to an ECM-bound state would be by affecting zinc coordination, thereby changing its Kd for zinc. We therefore tested if the EC_50_ of zinc is different under high (1.8 mM, the concentration in regular DMEM) and low (0.18 mM, the lowest concentration the cultured cells appeared normal) calcium. Under low calcium conditions, the addition of 5 μM zinc to the medium resulted in increased accumulation of ShhN in the ECM both of Shh-C199*-Ca_Free_ and Shh-C199* (Figure 3A). This indicates that Shh-C199*-Ca_Free_ is still able to mediate ECM accumulation, and supports the notion that calcium binding is not required for Shh distribution. E127 is located at the interface between the calcium and zinc-binding domains of Shh, and we tested if restauration of this residue in Shh-C199*-Ca_Free_ affects ECM localization but found little or no difference (Figure 3A). To better quantify the effect of calcium and zinc on Shh-C199* and Shh-C199*-Ca_Free_ in their ability to accumulate in the ECM we used ELISA directly on the de-cellularized ECM. Under low calcium conditions we found that the response to increasing zinc concentrations was similar between Shh-C199* and Shh-C199*-Ca_Free_. For both, the EC_50_ for zinc appeared to be around 0.1 μM. Instantiated in Figure 3A and quantified over multiple experiments in Figure 3B, it appears that Shh-C199*-Ca_Free_ is less efficient in entering the ECM than Shh-C199*. This effect was more profound in the presence of 1.8 mM calcium, and much more Shh-C199* was detected in the ECM than Shh-C199*-Ca_Free_ in the absence of added zinc. The addition of zinc had a bigger effect on Shh-C199*-Ca_Free_ than on Shh-C199*. These results indicate that Shh-C199*-Ca_Free_ behaves similarly in high and low calcium and resembles Shh-C199* under low calcium. Thus, whereas the behavior of Shh-C199* changes as a function of calcium, that of Shh-C199*-Ca_Free_ does not, indicating that binding of calcium to Shh alters its intrinsic properties as measured by its ECM association.

### Distribution of cholesterol-modified ShhNp in the ECM differs from cholesterol-unmodified ShhN but remains zinc sensitive

While ShhN could readily be detected in the ECM of Hek293t cells, processed ShhNp poorly accumulated in the ECM, possibly reflecting earlier bottlenecks, like processing (Lee et al., 1994), Dispatched1 activity (Ma et al., 2002) and re-internalization. As ShhN can be internalized via several Shh-binding proteins, (Incardona et al., 2000; McCarthy et al., 2002; Wilson and Chuang, 2010) we assessed if expression of Shh in cells lacking many of its binding partners altered ECM accumulation. Staining for Shh in the ECM of transfected fibroblasts lacking many Shh (co)-receptors (*Ptch1*_*LacZ/LacZ*_;*Ptch2_−/−_;Boc_−/−_*;*Cdo_−/−_*;*Gas1_−/−_*) showed that ShhN was present in small puncta that gave a cloudy appearance at lower magnifications (Figure 4A). 5 μM zinc increased the number of these small puncta, which could be quantified by measuring the fluorescence intensity across the entire image area (Figure 4B). In contrast, cholesterol-modified ShhNp was detected in larger puncta in a more restricted area. While the distribution of large ShhNp puncta with increasing zinc was largely unaffected, we detected an increase in small puncta with a wider distribution resembling ShhN distribution. The effects of zinc on Shh distribution in the presence of 1.8 mM calcium was much less pronounced, further supporting the finding that high calcium negatively affects the zinc-dependent activity of Shh. To assess if the observed effects require the zinc-coordination fold of Shh we tested ShhE177A, and - H181Y for their ability to associate with the ECM. Consistent with the biochemical observations using Shh-C199* (Figures 1, 2), we could barely visualize ShhE177A in the ECM. In contrast, ShhH181Y distribution into the ECM was indistinguishable from Shh, further indicating that this “butterfly version” of Shh is functional (Figure 4B).

**Figure 4:**
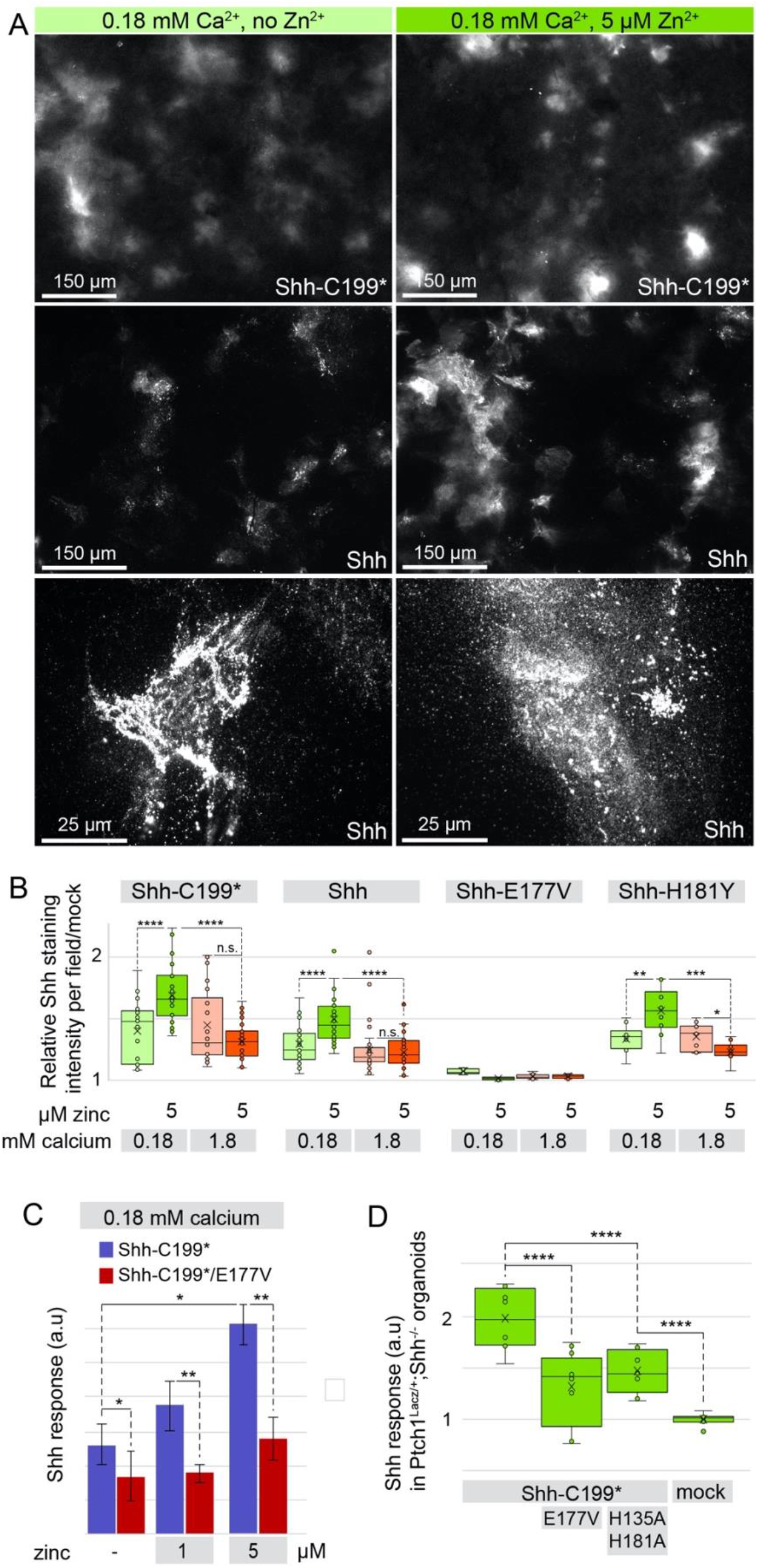
Cholesterol-modified ShhNp associates with the ECM in a zinc and peptidase-dependent manner. **A:** *In situ* staining with anti ShhN mAb5E1 of ECM deposited by *Ptch1_LacZ/LacZ_*;*Ptch2*_−/−_;*Boc*_−/−_;*Cdo_−/−_*;*Gas1_−/−_* cells that were transfected with *Shh* or *Shh-C199\** in the presence or absence of 5 μM zinc and presence of 0.18 mM calcium. **B:** Quantification of ECM-bound Shh shown in A. Box and whisker plots of mean fluorescence intensity per image of 10 microscope fields per experiment was measured in ImageJ and normalized to the ECM of mock transfected cells. n=3, \*\*\*\**p*<0.0001.). **C:** LightII cells co-cultured with *Shh-C199\** and *Shh-C199*/E177V* transfected *Ptch1*_LacZ/LacZ_;*Ptch2*_−/−_ fibroblasts in the presence of 0.18 calcium and the indicated zinc concentrations. Responses were normalized to Shh-C199* with 1.8 mM calcium and no zinc. Shown are means and standard errors of 3 independent experiments. \**p*<0.05, \*\**p*<0.01. **D:** HEK293t cells transfected with *Shh-C199\** and *Shh-C199*/H135A/H181A* were incorporated into spinal cord organoids otherwise consisting of *Ptch1*+/LacZ;*Shh*_−/−_ stem cells were assessed for LacZ induction. n=4, \*\*\*\**p*<0.0001.

To test if the increased accumulation of Shh in the ECM is correlated with the non-cell autonomous signaling efficacy, we co-cultured LightII cells (Taipale et al., 2002) with transfected fibroblasts. This allows us to strictly measure non-cell autonomous signaling, and we found that in the presence of increasing concentrations of zinc, non-cell autonomous signaling is enhanced. This effect of zinc was not observed in co-cultures with *Shh-C199*/E177V*-transfected fibroblasts, further indicating that zinc affects an intrinsic property of Shh that enhances non-cell autonomous signaling (Figure 4C). In a more stringent approach to assess signaling activity of Shh over long distances, we made aggregates of Shh-C199* transfected Hek293t cells and co-cultured them with organoids derived from mouse embryonic stem cells (mESCs). The Hh-response in the *Ptch1*_LacZ/+_*;Shh*_−/−_ organoids was increased approximately 2-fold in response to Shh-C199* (Figure 4D). The pathway response was significantly decreased in these organoids when co-cultured with Shh-C199*/E177V and Shh-C199*/H135A/H181A expressing aggregates, further supporting the idea that an intact zinc-coordination fold of Shh is necessary to mediate the release from Shh-producing cells and consequently, the facilitation of long-range signaling.

### The peptidase domain of a BacHh is unable to facilitate ECM association

In bacterial Hhs (BacHh), the residues of the zinc coordination fold described in this work are conserved, suggesting that the fold might exert a similar function. To test if this part of BacHh is sufficient to mediate the observed ECM accumulation, we made a construct coding for a chimeric protein consisting of the N-terminal 65 residues of Shh, the calcium and zinc binding domains of *Bradyrhizobium paxllaeri* BacHh (codon optimized for expression in mammalian cells), followed by an HA tag replacing the bacterial stop codon, followed by the last 10 residues of Shh up to G198 (Shh/BacHh_HA_, Figure 5A diagram). As a control, we positioned an HA tag at the same distance (10 residues) from the C-terminus of Shh-C199* (ShhHA-C199*). We found that ShhHA-C199* behaved indistinguishable from Shh-C199* and entered into the ECM and the medium in a zinc-dependent manner (Fig 5A). In contrast, although readily detected in the lysate, no Shh/BacHh_HA_ could be detected in the ECM or the medium (Figure 5A). To assess where in the cell the chimeric protein accumulates, we performed detergent-free live staining with an α-HA antibody prior to fixation of receptor-less *Ptch1*_*LacZ/LacZ*_;*Ptch2_−/−_;*Boc*_−/−_*;*Cdo_−/−_*;*Gas1_−/−_* fibroblasts transfected with the aforementioned constructs. We found no difference in staining between ShhHA-C199* and Shh/BacHh_HA_, indicating that Shh/BacHh_HA_, similarly to ShhHA-C199*, is being trafficked through the ER and Golgi apparatus to the cell membrane (Figure 5B). It is not, however, being released from the cell, indicating that the bacterial zinc-coordination domain is not sufficient for entry into the ECM. *Bradyrhizobium paxllaeri* BacHh presumably lacks the specificity for an ECM binding partner that is recognized by Shh.

**Figure 5:**
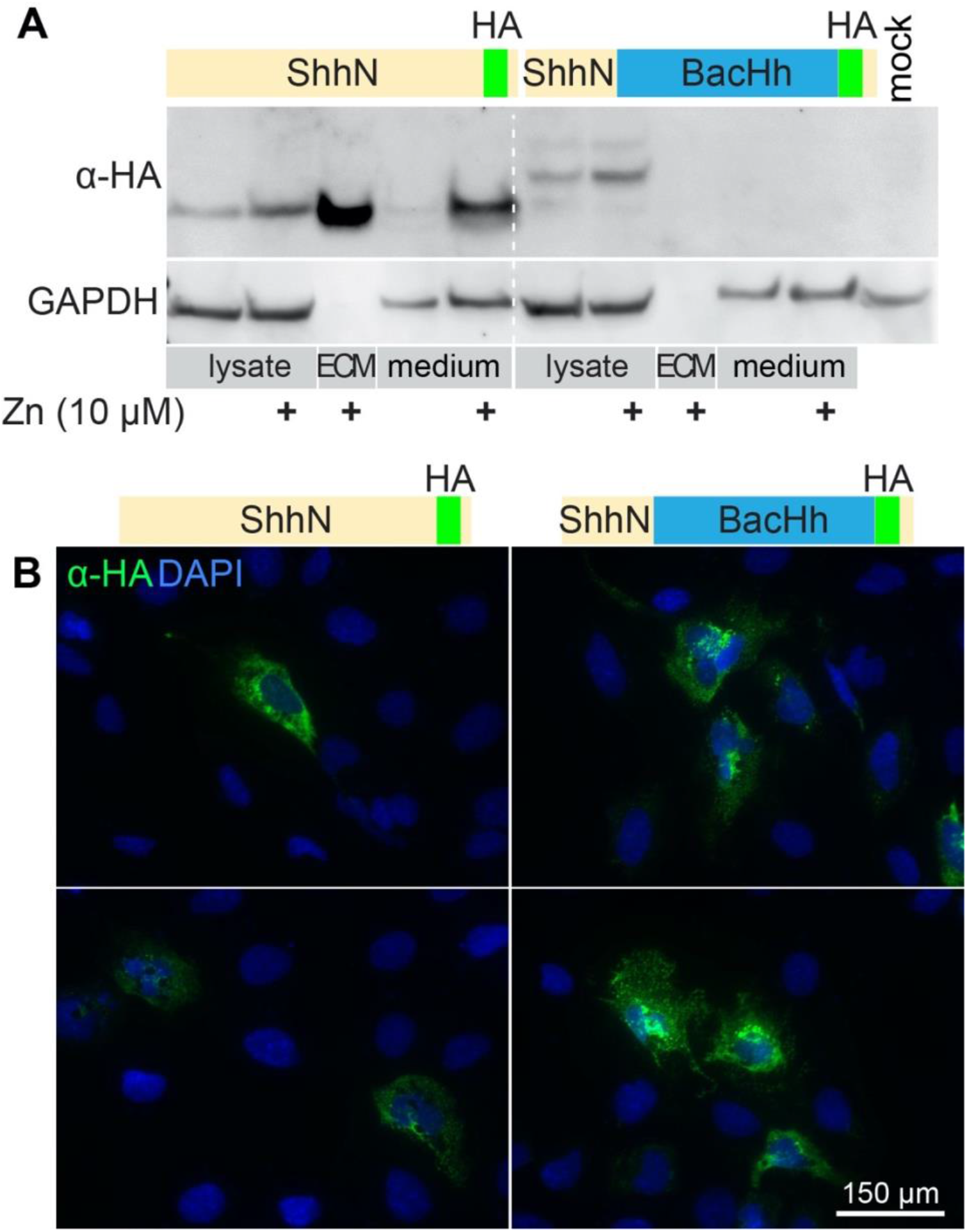
The zinc-coordination domain of BacHh is not sufficient for association with the ECM. **A:** Western Blot analysis of the lysate, ECM, and supernatant of *ShhN-HA* or *Shh-BacHh-HA* (diagrams) transfected HEK293t cultured in the indicated zinc concentrations. **B:** Detergent-free live staining with an α-HA antibody (3F10) of transfected *Ptch1_LacZ/LacZ_*;*Ptch2_−/−_;*Boc*_−/−_*;*Cdo_−/−_*;*Gas1_−/−_* cells. Nuclei were stained with DAPI.

## Discussion

Here, we provide evidence for a function of the zinc-coordination fold of Shh for the association of Shh to the ECM. Zinc is a potent agonist of this domain as it increases ECM association of Shh but not of zinc-coordination fold mutants. The observations that Shh-E177A is unable to mediate signaling from the notochord to the overlying neural tube (non-cell autonomously), and is more capable than Shh to induce the Hh response when expressed in the developing neural tube (likely cell-autonomously) (Himmelstein et al., 2017), provides in vivo evidence that the zinc-coordination fold of Shh is required for non-cell autonomous signaling, but not for the activation of the Hh response *pe se*. The initial experiments that demonstrated that E177 is dispensable for the activation of the Hh response is easily explained as this mutant ligand was added to the responding cells as a purified and soluble fraction (Fuse et al., 1999), thus bypassing the requirement for the function of the zinc-coordination fold that we propose. Distinct from the peptidase domain is the extreme N-terminal end of ShhN which binds to Ptch1 (Gong et al., 2018; Qi et al., 2018b; Qi et al., 2018a) and suffices to alter Ptch1 activity (Tukachinsky et al., 2016), further demonstrating the dispensability of the zinc-coordination fold to activate the Shh response in the responding cell.

### Do Bacterial Hhs and Shh share a peptidase activity?

Our observations indicate that Shh distribution away from the sites of synthesis and non-cell autonomous Shh signaling can be enhanced under low-calcium and high zinc conditions. The surprising sequence similarity between bacterial and mammalian Hedgehog proteins strongly suggest they have similar functions. The organization of bacterial genomes into operons helps in the assignment of possible functions of unknown proteins. The suggested role of BacHh (as a M15A peptidase (Rawlings and Barrett, 2013)) in the modification of the bacterial peptidoglycans is further supported by the observation that in *Mesorhizobium* and *Bradyrhizobium* the *BacHh* gene is surrounded by genes (likely constituting an operon) that code for proteases, including lysozyme, N-acetylmuramoyl-L-alanine amidase, a peptidoglycan endopeptidase (peptidase M23A), several Trypsin homologs (peptidase S1), a zinc Matrix Metalloprotease (MMP) homolog (peptidase M54), an endonuclease, peptidase S53, and possibly a Phytase (DUF3616). This complex of enzymes might be involved in bacterial feeding or scavenging. BacHhs in *Rhizobiacea* are not part of the core genome (González et al., 2019), as the majority of these bacteria do not carry *BacHh*, a further indication that BacHhs provide a niche-specific specialized function.

Possible mechanisms of catalysis of zinc peptidases have been elucidated with the help of structural models of enzyme-inhibitor complexes. Thermolysin is a well-studied zinc metallopeptidase structurally related to Shh (Hall et al., 1995; Rebollido-Rios et al., 2014). Shh and Thermolysin coordinate zinc via two histidine and an aspartic acid residue. A catalytic glutamic acid residue initiates catalysis (E177 in mouse Shh) by accepting a proton from water to form the nucleophilic hydroxide that attacks the carbonyl carbon, further stabilized by the coordinated zinc. With the two stacking histidine (or occasionally tyrosine) residues a pentacoordinate transition state is formed and resolved into the hydrolyzation products (Matthews, 2002; Tronrud et al., 1992).

M15 peptidases cleave peptidoglycans, the major component of the bacterial periplasmic space, and a major component of detritus. Peptidoglycans are analogs of proteoglycans that are common in extracellular matrix (ECM) of animals. Therefore, it is possible that functional conservation between BacHhs and Shh is reflected in the ability of Shh to cleave or modify proteoglycans, thus affecting the Shh response and/or distribution, independent of its binding to the canonical Hh receptors. Although any Shh antagonist could be a possible target for the putative Shh peptidase activity, the Hh-interacting protein (Hhip) is an unlikely candidate as it binds to Shh via the zinc ion, thereby replacing the catalytic water. This mode of binding is akin to that of a metalloprotease/inhibitor interaction (Bosanac et al., 2009), and thus likely to inhibit the putative catalytic function of Shh instead of being a substrate. The targets for penicillin and related antibiotics are peptidoglycan peptidases, leaving open the development of nM drugs that inactivate the peptidase activity of Shh, and thus be powerful inhibitors of non-autonomous Shh signaling that underlies several cancers.

### Are Hhs proteoglycan peptidases?

Hedglings and Hedglets are related to Hhs, and the conserved domains possibly homologous. All animals that have Hedglet also have a *Hh* gene and it is thus plausible that Hedglets are derived from Hh. Hhs are not found in any eukaryote except *cnidarians* and bilaterians. The distribution of Hedgling and Hhs only overlaps in *cnidarians*, but Hedgling can also be found in sponges and choanoflagellates (Figure S1). This suggests two evolutionary events giving rise to these proteins; one occurring in a *Choanoflagellate* ancestor that originated the gene coding for Hedgling, and an independent event in a *Cnidarian* ancestor that gave rise to modern Hh. The absence of both Hedgling and Hh from algae, plants, fungi, in addition to almost all unicellular eukaryotes makes it unlikely that both Hh and Hedgling linearly evolved from a BacHh protein that could have been present in the Ur-eukaryote, but more likely are products of more recent gene transfers from bacteria. The distribution among eukaryotes of Glypicans and Hs is overlapping, and both are first observed in *Cnidarians* and present in all bilaterians. A more recent evolutionary relationship between BacHh and Hhs is further supported by the observation that the C-terminal residue of many BacHhs perfectly aligns with the exon 2 splice donor site in *Hh* genes, thereby providing a parsimonious explanation how a *BacHh* gene was incorporated in a eukaryotic genome giving rise to *Hh*. This is in contrast to the much less conserved Hh domain in Hedgling that is encoded within a single large exon. Given the central role of Gpcs in the distribution of and response to Hhs (including Shh), Glypicans and Hhs might have co-evolved possibly as a peptidase/substrate combination, co-opting the peptidoglycan activity of BacHhs to cleave the proteoglycan Glypican. Hh-like bacterial peptidases (M15A) are predicted to be carboxy(trans)peptidases, cleaving adjacent to the D-ala that is linked to the murein glycans (Bochtler et al., 2004; Vollmer and Bertsche, 2008). By analogy, Shh might cleave an unusually modified C-terminal residue. It is intriguing that the C-termini of Glypicans are linked to the GPI anchor that restricts them to the cell surface (Filmus et al., 1995). Solubilization of Shh-sequestering Gpcs by GPI removal would elegantly reconcile the observed peptidase-dependent entry of Shh into the ECM with the important effects of Gpcs on Shh signaling and distribution.

It is perhaps unfortunate that Hh was discovered in *Drosophila,* as of all animals sequenced, only Hh in *Drosophilids* is divergent for two of the three residues that coordinate zinc and lacks the critical E177 equivalent. The predicted lack of peptidase activity in *Drosophilid* Hhs is remarkable and further supports the observation that the putative peptidase activity is not required for the Hh/receptor interaction. Perhaps stricter reliance on cytonemes in *Drosophila* that detect Hh at its source (Huang et al., 2019) renders the ancestral peptidase activity obsolete. Nevertheless, this loss of the putative peptidase activity is unique to *Drosophilids*, as all other (sequenced) animals retain the typical zinc coordination motif and the associated E177 equivalent that are required for catalytic activity. Based on the loss-of-function of several mutants, this intrinsic property is likely a zinc metallopeptidase activity, just like the bacterial counterparts of Shh. Still, the observation that substitution of the Shh calcium/zinc domains with those of BacHh results in a protein that does not enter the ECM, indicates that their substrates are not interchangeable.

Although mutations of the central zinc coordinating triad are unstable and thus cannot be easily assessed for loss of peptidase function, mutations of several other catalytically important residues (E177, H135 and H181) are not destabilized and show a loss in the ability of Shh to enter the ECM. Together with the observation by Himmelstein and colleagues (Himmelstein et al., 2017) that ShhE177A cannot signal from the notochord to the overlying neural plate strongly supports the idea that a Shh-associated peptidase activity is required for non-cell autonomous signaling by promoting its distribution away from the source cells.

## Material and Methods

### Sequence analysis

Bacterial Hedgehogs, Hedglings and Hedglets were identified via protein-protein BLAST (NCBI) and HMMER (ensemble) searches (Gish and States, 1993) using the peptide sequence of the Shh N-terminal domain as the initial query sequence. Conserved sequences were manually curated to contain only the calcium and zinc coordination domains (around 105 residues). Sequences (supplemental file) were aligned in Clustal Omega (EMBL-EBI). An average distance tree and a PCA plot were generated in Jalview (Waterhouse et al., 2009), using the BLOSUM62 algorithm. Visualizations of the ShhN structure were generated in UCSF Chimera using Protein Database (PDB) ID 3D1M (McLellan et al., 2008).

### Materials

MG-132 and Concanamycin A were from Calbiochem, Chloroquine and ZnCl2 from Sigma, CaCl_2_ from Fisher Scientific, and Dynasore from Abcam.

### Cell culture

*Ptch1_−/−_;Ptch2_−/−_* fibroblasts were derived from mouse embryonic stem cells and are described elsewhere (Casillas and Roelink, 2018). All cells were cultured in DMEM (Invitrogen) supplemented with 10% FBS (Atlas Biologicals). Mouse embryonic stem cells were cultured in DMEM (Invitrogen) supplemented with 20% FBS (Atlas Biologicals), 2 mM L-Glutamine (Gibco), 1X MEM non-essential amino acids (Gibco), 0.001% 2-Mercaptoethanol (Gibco), and 0.001% Leukemia Inhibitory Factor (LIF). Cells were transfected using Lipofectamine2000 reagent (Invitrogen) according to the manufacturer’s protocol.

### DNA constructs

The following mutations were created via site-directed mutagenesis: *Shh-C199*/E177V, Shh-C199*/H135A, Shh-C199*/H135Y, Shh-C199*/H181A, Shh-C199*/H181Y, Shh-C199*/H135A/H181A, Shh-C199*/H135Y/H181Y, Shh-C199*/E90A/E91D/D96A/D130N/D132L, Shh-C199*/E90A/E91D/D96A/E127A/D130N/D132L*. *Bradyrhizobium paxllaer*i *BacHh* (EnsemblBacteria: LMTR21_38280, NCBI: WP_065756078.1) was codon optimized for eukaryotic expression using the IDT DNA Codon Optimization Tool, ordered as a gBlocks gene fragment from IDT DNA, and cloned into pcDNA3.1(+). Both the *Shh-C199\** vector backbone including the Shh N- and C-terminus as well as the calcium and zinc coordination domain of *Bradyrhizobium paxllaeri BacHh* were PCR amplified, separated on a 1% agarose gel, and extracted with MinElute columns (QIAGEN). The fragments were cut with *BsaI* and ligated with T4 DNA ligase according to the Golden Gate cloning protocol (New England Biolabs).

### Immunostaining

*Ptch1_−/−_;Ptch2_−/−_* fibroblasts were plated on 12mm glass cover slips and transfected with *Shh-C199\** the following day and subsequently allowed to recover for 24h. The transfected cells were then incubated for 24h in serum-free DMEM containing varying concentrations of CaCl_2_ or ZnCl_2_, the cells were detached from the cover slip with PBS. The cover slips were washed with PBS at least 5 times and blocked with 10% heat-inactivated goat serum in PBS with 0.1% TritonX (PBS-T). Mouse α-Shh (5E1, Developmental Studies Hybridoma Bank) was used at 1:30 in blocking solution and goat α-mouse Alexa568 secondary antibody (Invitrogen) at 1:1,000 in blocking solution. Shh distribution was visualized with a Zeiss Observer at 10x and 63x magnification.

For live staining, transfected cells were incubated in serum-free medium for 20h. An α-HA antibody 3F10 (Sigma) was added for another 4 hours (1:1,000) before cells were fixed with 4% PFA in PBS and blocked in 10% heat-inactivated goat serum in PBS. A goat α-rat Alexa488 secondary antibody (Invitrogen) was used at 1:1,000 in blocking solution and nuclei stained with DAPI.

### Western Blot/SYPRO ruby staining

Hek293t cells were plated in 12 well plates and transfected with Shh mutants as indicated the next day. 24h after transfection, the medium was switched to serum free DMEM with the indicated calcium and zinc concentrations overnight. Cells were then detached from the plate with PBS and lysed in a microcentrifuge tube with RIPA buffer (150 mM NaCl, 50 mM Tris-HCl, 1% Igepal, 0.5% Sodium Deoxycholate, and protease inhibitors). The lysate was incubated for 30 min on ice and cleared by centrifugation. For isolation of ECM-bound Shh, the decellularized tissue culture dish was washed with PBS and deionized water at least 5 times and scraped with a cell scraper and 5X SDS sample buffer heated to 95oC, as described (Hellewell et al., 2017). A fifth of the sample was run on a 12% SDS-PAGE gel and transferred to a 0.45μ nitrocellulose membrane. Membranes were blocked in 5% milk in Tris-buffered saline with 0.1% Tween-20 (TBS-T) and incubated with a polyclonal rabbit α-Shh antibody (H2, 1:10,000) (Roelink et al., 1995) in blocking solution, followed by incubation with a goat α-rabbit Alexa647 secondary antibody (Invitrogen, 1:10,000) in blocking solution. GAPDH was used as a loading control (Rabbit α-GAPDH, 14C10, Cell Signaling Technologies). Western Blots were visualized with a ChemiDoc visualization system (Bio-Rad).

Alternatively, the SDS-PAGE gel was stained with SYPRO-Ruby gel stain (Thermo-Fisher) according to the manufacturer’s instructions and visualized with a ChemiDoc visualization system (BioRad).

### ELISA

Hek293t cells were plated in 96 well plates and transfected with *Shh-C199\** and *Shh-C199*/E90A/E91D/D96A/E127A/D130N/D132L* in triplicates the next day. 24 h after transfection, the medium was replaced with DMEM containing 0.18 mM or 1.8 mM Calcium and Zinc concentrations ranging from 0.001 to 1 μM for 48 h. The cells were removed from the plate with PBS and deionized water. The plates were blocked with PBS + 5% heat-inactivated goat serum, incubated with mAB5E1, followed by an HRP conjugated α-mouse secondary antibody (Invitrogen). Western-Lightning Plus-ECL (Perkin Elmer) was added to the wells and luminescence was measured in a Wallac Victor3 plate reader (Perkin Elmer).

### Non-cell autonomous signaling assay

*Ptch1_−/−_;Ptch2_−/−_* fibroblasts were plated in 24 well plates and transfected with the indicated *Shh-C199\** variants the next day. 24h after transfection, cells were washed with PBS once and LightII reporter cells (Taipale et al., 2000) were added. As soon as cells were adherent, the medium was switched to DMEM containing the indicated zinc and calcium concentrations. 48h later, the cells were lysed and luciferase activity was measured using the Dual Luciferase Reporter Assay System (Promega). Firefly luciferase measurements were normalized against Renilla luciferase measurements for each technical replicate to control for differences in cell growth. Firefly/Renilla luciferase values were then normalized to the mock control average for each experiment.

For the long-range signaling assay, Hek293t cells were plated in 12 well plates and transfected with the indicated Shh-C199* constructs the next day. 24h after transfection, cells were washed off with PBS and collected in a conical tube. Aggregates were allowed to form for 48h in DFNB medium in non-treated tissue culture plates rotated at 1 Hz. Similarly, four times as many mESCs as Hek293t cells were aggregated in DFNB medium for 48h rotated at 1 Hz. Hek293t aggregates and 2 μM Retinoic Acid (RA) were added to the mESC organoids for another 48h. The cells were then collected, washed once in PBS, and lysed in 100 mM potassium phosphate, pH 7.8, 0.2% Triton X-100. The lysates were cleared by centrifugation and 20 μl analyzed in triplicates for Ptch1:LacZ expression using the Galacto-Light chemiluminescent kit (Applied Biosciences). Lysates were normalized for total protein using the Bradford reagent (BioRad).

### Genome editing

*Ptch1*_LacZ/LacZ_;*Ptch2*_−/−_;*Boc*_−/−_;*Cdo*_−/−_;*Gas1*_−/−_*;Shh_−/−_* were derived from *Ptch1*_LacZ/LacZ_;*Ptch2*_−/−_-*;Shh_−/−_* cells (Roberts et al., 2016). TALEN constructs targeting the first exon of mouse *Cdo* and *Gas1* were designed and cloned into the pCTIG expression vectors containing IRES puromycin and IRES hygromycin selectable markers (Cermak et al., 2011). The following repeat variable domain sequences were generated: Cdo, 5’ TALEN: NN HD NI NG HD HD NI NN NI HD HD NG HD NN NN; 3’ TALEN: HD NI HD NI NI NN NI NI HD NI NG NI HD NI NN; Gas1, 5’ TALEN: NN NI NN NN NI HD NN HD HD HD NI NG NN HD HD; 3’ TALEN: NN NN NI NI NI NI NN NG NG NG NN NG HD HD NN NI. Two CRISPR constructs targeting a double strand break flanking the first exon of mouse Boc were cloned into pSpCas9 vector with an IRES puromycin selectable marker (Ran et al., 2013). The Boc CRISPRs targeted the following forward genomic sequences (PAM sequences underlined): Upstream of first exon 5’ <u>CCT</u>GTCCTCGCTGTTGGTCCCTA 3’; Downstream of first exon 5’ <u>CCC</u>ACAGACTCGCTGAAGAGCTC 3’. *Ptch1_LacZ/LacZ_;Ptch2_−/−_;Shh_−/−_* mouse embryonic stem cells (Roberts et al., 2016) were transfected with 6 genome editing plasmid. One day after transfection, ES medium with 100 ug/mL hygromycin and 0.5 ug/mL puromycin was added for 4 days. Surviving mESC colonies were isolated, expanded and genotyped by sequence PCR products spanning TALEN and CRISPR-binding sites. PCR screening was performed on cell lysates using primers flanking the TALEN or CRISPR binding sites for the *Boc*, *Cdo*, and *Gas1* loci. *Boc*, (5’) CATCTAACAGCGTTGTCCAACAATG and (3’) CAAGGTGGTATTGTCCGGATC; *Cdo*, (5’) CACTTCAGTGTGATCTCCAG and (3’) CCTTGAACTCACAGAGATTCG; Gas1, (5’) ATGCCAGAGCTGCGAAGTGCTA and (3’) AGCGCCTGCCAGCAGATGAG. PCR products were sequenced to confirm allele sequences. A *Ptch1*_LacZ/LacZ_;*Ptch2*_−/−_;*Boc*_−/−_;*Cdo*_−/−_;*Gas1*_−/−_*;Shh_−/−_* mESC clone was identified harboring a 50 bp deletion in Cdo exon 1, a heteroallelic 480 bp insertion and a 200 bp deletion in Gas1 exon1 resulting in a premature stop codon in the reading frame, and a 450 bp deletion of Boc exon 1. These cells were transfected with *LargeT* and *myc*, and deprived of Lif to generate immortalized fibroblasts.

## Data Analysis

Single Factor ANOVA was used to analyze more than two conditions, followed by a Student’s *t*-test with a two-tailed distribution assuming unequal variance comparing two conditions. \**p*<0.05, \*\**p*<0.01, \*\*\**p*<0.001, \*\*\*\**p*<0.0001.

## Acknowledgements

This work was supported by NIH grant 1R01GM117090 to HR. We thank Drs. N. King and D. Rokhsar for discussions on Hh evolution, and Dr. C. Casillas for help with the receptorless cell line.

## Author contributions

All experiments were performed by CJ. Experiments were designed by CJ and HR. The manuscript was written by CJ and HR.

---

**Figure S1:**
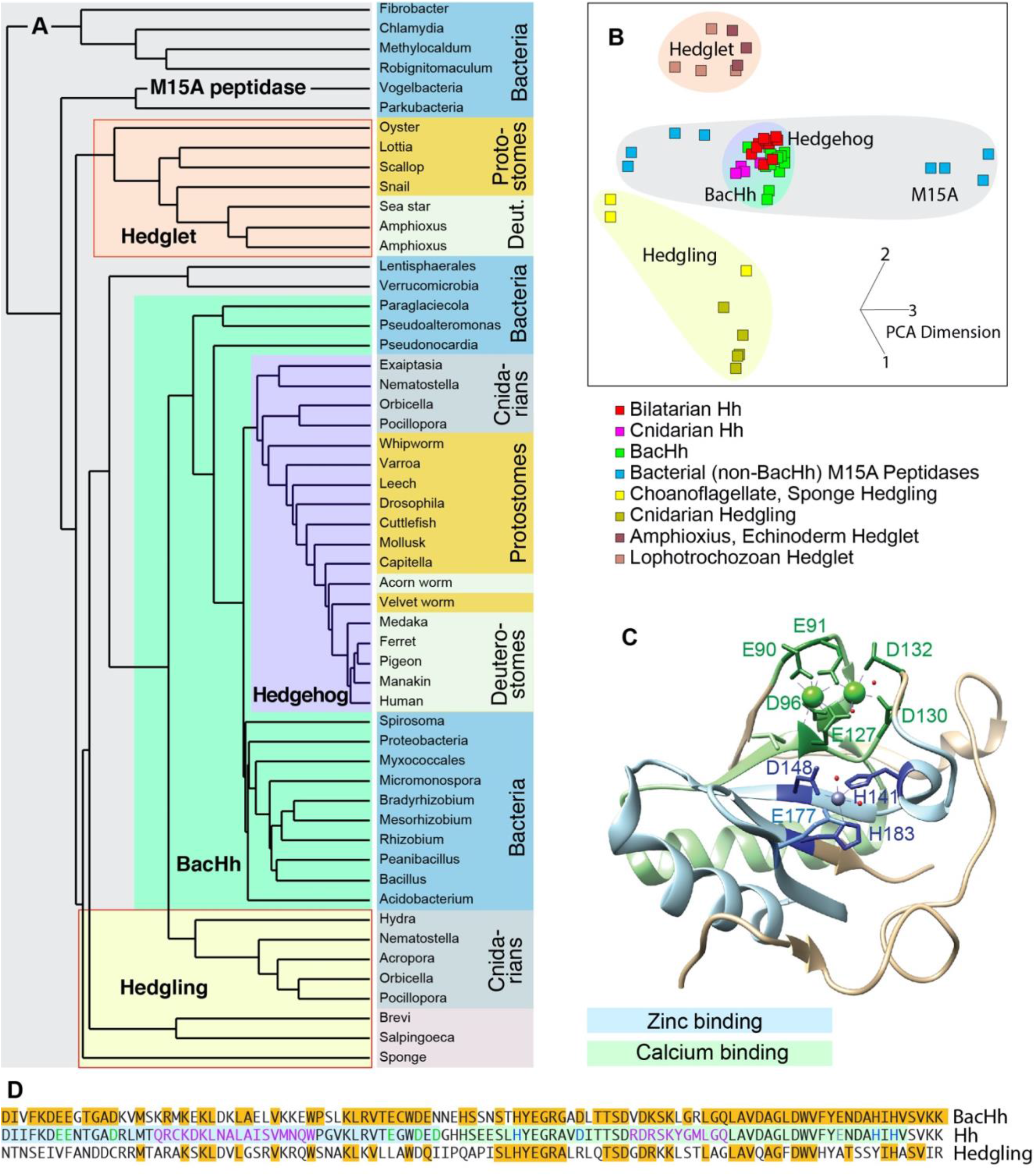
Sequence similarities between Hh, Hedgling, and Hedglet. **A:** a similarity tree of Hh sequences encompassing the calcium and zinc coordinating domains. All Hhs are closely related and root from within the BacHhs. Hedglings (present in Choanoflagellates, Sponges, and Cnidarians) and Hedglets (present in Lophotrochozoans, Amphioxus and Echinoderms) form outgroups, but all are related to bacterial M15 peptidases. Organisms in the same phylum or clade are color coded, and common names of the organisms are used. Sequences and accession numbers can be found as a supplemental file. Hedglings and Hedglets (red borders) are predicted to be unable to mediate catalysis. **B:** PCA plot of the tree presented in A. **C:** Structure of Shh with salient residues and domains indicated. **D:** Sequence lineup of Mesorhizobium BacHh, human Dhh, and Choanoflagellate Hedgling. Identical residues are indicated in amber. Salient residues are color coded and color coordinated with C.

**Figure S2:**
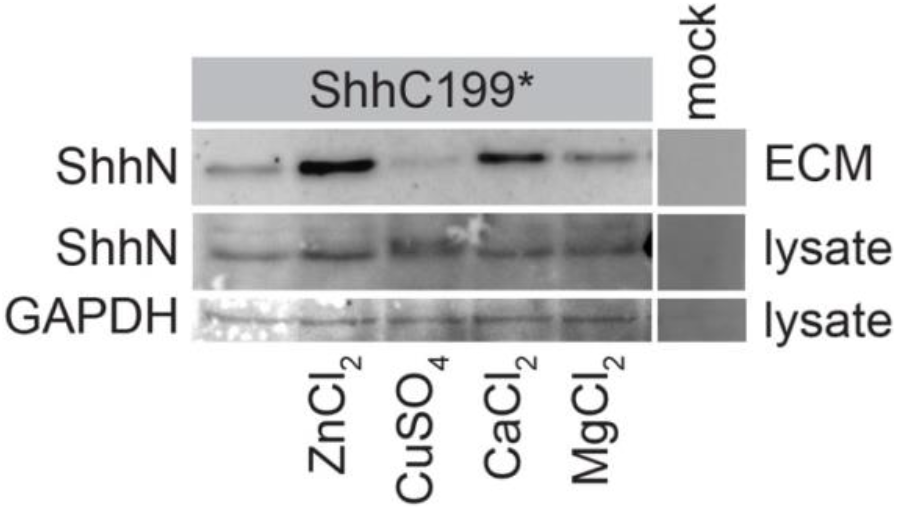
Association of Shh with the ECM is enhanced specifically in response to zinc. Western blot analysis of the lysate and ECM of Hek293t cells transfected with *Shh-C199\** and cultured in medium containing 0.18 mM calcium and 5 μM zinc, 5 μM copper, 1.8 mM magnesium or 1.8 mM calcium as indicated.

## Sequence supplement

~~~
>Monosiga brevicollis, XP_001749037.1, 354-458
GLSFAPSSGNMYPNVVSARMASRLKVLANLVPRVFGESAAVLVLDAYRAAPLVAAEATLHNTGRAALLTVINVTASLDDELA
ALASVCADAGFDYVLYNSSAAIY
>Human, BAA33523.2, 89-193
IIFKDEENTGADRLMTQRCKDRLNSLAISVMNQWPGVKLRVTEGWDEDGHHSEESLHYEGRAVDITTSDRDRNKYGLLARLA
VEAGFDWVYYESKAHVHCSVKSE
>Mesorhizobium sp. L103C131B0, ESZ55121.1, 173-277
IVFKDEENTGADRMMTPRLKSKLDSLANVVASEWPGAKLRVTEAWDEDNEHADASLHYEGRAADLTTNPVDGAKLGRLARLA
VDAGCDWVFFEDSSHIHVSVKAG
>Micromonospora sp. HK10, WP_082159544.1, 1063-1166
IVFKDEEKTDADRMMTPRLRDMVNELAALVVKEWPGKKLRVTEGWDENNEHTAESTHYEGRAVDMTVSDLDAAKLGRLARLA
VDAGFDWVFYENALHVHASVKK
>Nematostella1 vectensis XP_001635678.1, 84-187
IVFKDEERTGADRLMSKRCREKLRNLATKVKQKWKGVKLRVTEAWDEDGQHSLDSLHYEGRAVDISTSDKDPKKLPDLGSLA
VDAGFDWVYYDRRSSIHASVRS
>Nematostella2 vectensis, ABX84114.1, 69-171
EVVFENDDCRRTTARAKSKLDVLASRVRQEWAGRKLKVIKAWTDQRTAQDPASLHYEGRALRLQLDNNDRSMLSRLAGLALA
SGFDWVSYPLNSDYIHASVIRA
>Pseudoalteromonas piratica, WP_040135141.1, 60-162
PVFKFEEGNFTDVQASEKLCAAIMDLNKLVMKEWPGKTLRVTEAYDQDGEHAKFSLHNEGRAADMTVSDRDLKKLGRLGFLA
TKAGFSWVYYEHNHIHASVKR
>Spirosoma aerolatum, WP_080055297.1, 211-317
VVFKNEEGDGSDRMMTPVLKTHVDRLADLVRSEWGAGVSLRVTEAWDDTGEHSSSHSLHYEGRAVDLTTSDLDKSKLGRLGR
LAVDAGFNWVYYENLLHIHASVTKA
>Varroa destructor, XP_022667503.1, 83-186
IRFLDDEGTGADRIMTQRCRDKLDTLAVSVMTQWPGVKLRVIESWDEYSHHKSGSLHYEGRAVDFTTDDRHQAKYGMLARLA
VEAGFDWVYYETKRHVHASVKP
>Drosophila melanogaster, NP_001034065.1, 144-247
ILFRDEEGTGADRLMSKRCKEKLNVLAYSVMNEWPGIRLLVTESWDEDYHHGQESLHYEGRAVTIATSDRDQSKYGMLARLA
VEAGFDWVSYVSRRHIYCSVKS
>Crocodylus porosus,XP_019386078.1, 87-190
IIFKDEENTGADRLMTQRCKDKLNALAISVMNQWPGVKLRVTEGWDEDGHHSEESLHYEGRAVDITTSDRDRSKYGMLARLA
VEAGFDWVYYESKAHIHCSVKA
>Amphimedon queenslandica, ABX90059.1, 74-180
SSQATYLHFASSDCRIMSSRLYTRLSSLAEAYYWRYHIKILVLKAWTPYPDYSLDNTSLHYEGRSVRIHVTSRNVTRLLKMA
VSAGFDWVMYDKKGYARMSVIPDAC
>Salpingoeca rosetta, XP_004997926.1, 540-646
TVKPDPPTSNGDPSVMSKRLRRHITTLASVVRGVFGDDAYVRVLEAYVEPPADISKASLHNVGRAARITIEGVPDDFASDRL
GVLGGLAVEAGFDYVAYTSRDSLYV
>Orbicella1 faveolata, XP_020616832.1, 68-170
IVFANDDCRRMTARAKSKLDVLGSRVKRQWSNAKLKVLLAWTDQIIPQAPISLHYEGRALRLQTSDGDRKKLSTLAGLAVQA
GFDWVHYATSSYIHASVIRDV
>Pocillopora1 damicornis, XP_027055961.1, 66-167
EIDFANDDCRRMTARAKSKLDVLGSTVRRQWSNVKLKVTLAWTDQIMPQAPISLHYEGRAVRLQTSDGDTGKLSTLAGLAVQ
AGFDWVHYATNSYIHASVIR
>Hydra vulgaris, XP_004209904.1, 72-174
IDFQTEDSRLMTSRAKQKIDTLAGLVTTRFGKNMKVNVLKAWTDVVEKEDKLSLHYEGRAFLIRASNNDKKLLSDLMVLARE
AGFDWVYYKNEDSIYLSVIPD
>Orbicella2 faveolata, XP_020632016.1, 84-183
IIFRDEEGTGADRLMSKRCKEKLTTLAGLVKGEWPSVKLVVTEAWDEQDQHSPNSLHYEGRAVDLRLSDRDKTKIGLLGRLA
VEAGFDWVLYESRSHIHA
>Pocillopora2 damicornis, XP_027044648.1 88-190
IIFKDEEGTGADRLMSKRCQDKLNTLADLVRRQWPTVKLVVTEAWDEQGQHSENSLHYEGRAVDLRLSDKDRTKIGYLGRLA
VDAGFDWVYYQKRTHIHASVR
>Exaiptasia pallida, XP_020892909.1, 87-190
IVFKDEEGTGADRIMSKRLREKLRILAKKVKEKWRGSTRLRVIEAWDEDGTHSAHSLHYEGRAVDITTSDLDKQKYPELGRL
AVEAGFDWVFYESQEHIHASVY
>Acidobacteria bacterium, PYS76727.1, 46-149
IVFKDEEHTGDDRMMTSRLSARVDDLAARVKREFPGLKLRITEAWDDSTIHAPTSRHLEGRAVDITTSDVDHHKLGRLAGLA
VEAGFDWVFFENDLHVHASVKK
>Bacillus pseudomycoides, WP_098188151.1, 989-1092
IQFKDEEGTGADFLMTSRLSDKLNTLAILVNQEWPNIKLRVTEAWDEDNEHSSGSTHYEGRAADITTSDRDGNKLGRLAQLA
VDAGFDWVYYENKYHIHVSVKK
>Proteobacteria bacterium, PZN23661.1, 96-197
IVFKDEEATGADLLMTPRLRTRLHELARLVTREWPGVRLRVTEAWDEDSEHGENSIHYEGRAVDVTTSDRDRRKLGRLAGLA
IQAGFDWVSHERDHVHASVR
>Vogelbacteria bacterium, OHA60126.1, 199-302
ANGYSGINGPGRTDRVHRDVAEATVWVQQNLNSDHNLSTQITAAHTEGVGHSAGSEHYEGRAVDIQPTGGNVTSSNLNIIAD
YCRQAGFTYVLVENRHVHCDAR
>Methylocaldum marinum, WP_119628113.1, 46-148
KSQPPIAVKKGAILAGLDRRMYFALQKARRVWSRYGKLLVVTSGLDGRHKKGSLHYVGLAVDLRSRYFAPSTRRTVTRELRR
NLGDEFQVIDEKHHIHVEFDP
>Fibrobacter sp., WP_143394061.1, 26-138
NLLIKRVQLKTGVYTGKLDAAMDSAGLVVVAEYHKVMGDSYRPTITSANDYGKHARRSKHYENKALDFRISDVPRNKRSQLV
ASIRQALGKRFNVFWEEKNTANEHLHIELKE
>Chlamydia trachomatis, CRH64334.1, 1-102
MLQFKNNVRLSGVQEEILFIIDRIQRYFEVRLPKRDFVITSLTDGAHMKGSLHPKGLALDMRSRTLDKKEIEYFVTWFRKNF
EKSYDLVVEIDHIHIEYDPK
>Parcubacteria group bacterium, PSO44215.1, 232-331
ASGFRGIAGPGRTGKVKQPWVEKTKQIQEICENRYGGRPFQVTAACTYGVGHSNDSTHYRGEAVDLDPVDATNQQVISCVKE
AGGVPYYLDEDSHIHISS
>Robiginitomaculum sp., PHQ68463.1, 158-262
DENDIDIKEGADISDLTDDMTDTFDDISEAWADEAPGVTPVITSGGDGTHSTNSLHYDGNAVDLRTNNLTQAQTTTVASALS
TSLGSDYDVVVESDHIHVEYDPG
>Lentisphaerales bacterium, TFH13511.1, 91-194
WESDHDGENDEDDHLMHRGVQPLLNQLEKAVASCGAALKVHDASRPSGGGHCATSLHKEGRALDLTADGLTLEDLAKLCWVA
GFDWVFNENKRGAEHVHCSSRA
>Paraglaciecola hydrolytica, WP_068382217.1, 63-166
VVIKFEEGDCSDSKVTKNLKKTIFKLVELIDQEWEGERKLRITEAWDNNAEHTKYSLHNEGRAADITTDDRDTKKLSKLACL
AMAAGFSWVKLEKDHVHASVPR
>Rhizobium leguminosarum, WP_130783679.1, 310-413
IVFKDEEGTGADRMMSARLRDGLDRLAAQVGIEWPDVKLRVTEAWDENNEHHGASLHYEGRAADLTTSPRDGDKLGRLGKLA
VDAGLDWVFFENSAHIHVSVKR
>Verrucomicrobia bacterium, HCF95878.1, 132-232
ESDHDGDWDTENDHLVHRDILPALIRLNALVLQEGATLKIQDAYREEGIHAPASLHREGRALDLTADGMSLARLAQLAVQAG
FDWVYYESPKGGGAHIHAS
>Myxococcales bacterium, RYZ03269.1, 109-212
IVFKDEERNRSDRFMTPRLRRSLVQLSKLVSQTWPKVDLRVTEAWDDRREHGAGSVHYEGRAADITTSDQDPAKLGTLAALA
VKAGFDWVFYENATHVHVSVKR
>Branchiostoma1 floridae, XP_002599309.1, 305-414
RMLGSSLDDRCADRVMSKALLDHLRTVQRMVQDEFSGVKLKVLEAWDEPHAGATTGDHPAGSLHYEGRAAKLTLSDGDAAKL
PRLAAFCICDGAGYVENKGDHILVAVQK
>Pseudonocardia dioxanivorans, WP_103381118.1, 106-208
VVKDEEGSGADRMMTPRLAELVGVLAAHVAQAFPGRRLRLTEAWDPDGEHSHSSLHYEGRAADLTVDDRDRAKLGRLAALAV
QTGFDWVLHENDHVHVSVRAG
>Branchiostoma2 belcheri, XP_019614930.1, 311-413
DRCADRVMTKSMLDLLRKVQKMVKDEFTGVKLKVLEAWDEPHAGATEGDQPAESLHFEGRAAKLTLTDGDTSKLPQLAKNAI
CAGANFVEHKGDHIFVAVRKQ
>Acanthaster planci, XP_022111291.1, 310-422
HMKGFALNSRCADRTMSARLMATLKTLGKLVSIEWPGVKLLVLEAWDEAHEGSTYTDGDQPAGSLHYEGRAAKLSLSDGDTS
KFSRLAGLATCAAADYVEHNGDHIFVAAKKQ
>Sepia bandensis, ALM01450.1, 32-143
IVFRDEESNNEDRMMSKRCKDKLNTLAIAVMNEWPGVKLRVTEAWDTEGHHAPTSLHYEGRAVDITTSDRERSRYGMLARLA
VEAGFDWVYYESRSHIHCSVR
>Lottia gigantea, XP_009064322.1, 299-407
EKPLGNSLNQRCAARLMSQRMYNVLISLQKLVRANGDKLKVEQAFDEKYAGHVADFDATSLYTEGRLVKVTRSVNPSLANYK
KLTQWAICSKADFVQNNGDHVLIGVKK
>Crassostrea virginica, XP_022317995.1, 275-379
YPGNYLPNRCAVRVMSPRLFNVLVNLKAYASDANLGGPGGKITVEEAWDGGADPSSLRSEGRMIKVKLSAGNTAANLGKLAQ
LAICAKADHVSNMGTHLLLSVKK
>Mizuhopecten yessoensis, XXP_021349176.1, 304-415
GIVGSALSKRCAARTMSYRMYKVINTLQKFVRHNMTLTDKLKVLKAWDEPYADATTGDTSYSRLHTEGRAVVVQLVSSNTAS
NLEELSHFAICAGADFISHKGDKLEIAVKK
>Bradyrhizobium sp. WSM4349, XWP_018460114.1, 210-314
SIVFKDEEGTGADRMMSTRMQAKLDALASLVSAEWPGVKLRVTEAWDENDEHSPTALHYEGRAADITTQPPDGAKLGRLARL
AVNAGCDWVFYEDTNHVHVSVKK
>Paenibacillus sp. CAA11, XWP_108465644.1, 1027-1131
DIVFKDEEGTGADKVMSKRMKEKLDKLAELVKKEWPSLKLRVTECWDENNEHSSNSTHYEGRGADLTTSDVDKSKLGRLGQL
AVDAGLDWVFYENDAHIHVSVKK
>Columba livia, XPKK32334.1, 86-190
DIIFKDEENTGADRLMTQRCKDKLNALAISVMNQWPGVKLRVTEGWDEDGHHSEESLHYEGRAVDITTSDRDRSKYGMLARL
AVEAGFDWVYYESKAHIHCSVKA
>Ptychodera flava, XBAR45718.1, 88-192
DIIFKDEEGTGADRLMTQRCKDKLNSLAILVMNQWEGIQLRVTEGWDEDGHHAENSLHYEGRAVDITTSDRDKKKYGMLARL
AVQAGFDWVYFESKSHVHCSVRS
>Antalis entails, XAPD15681.1, 86-190
DVIFKDEEGTGADRMMSKTCRDKLDTLAIFVMNQWTGVKLRVTEAWDEEHHHAKDSLHYEGRAVDVTTSDRDRSKYGMLARL
AVNAGFDWVYYESRAHIHCSVNS
>Helobdella robusta, XAAM70491.1, 270-374
NIIFQNSEGTGADRVMSKRCSDKLNNLASLTMEQWPGVRLRVVEAWDEDETHPEDSLHYEGRAVDVTTSDKDKSKYGMLARL
AVEAGFDWVHYEYRSHIHCSVKS
>Oryzias melastigma, XACL81248.1, 46-150
DIIFKDEENTGADRLMTQRCKDKLNSLAISVMNQWPGVKLRVTEGWDEDGHHFEESLHYEGRAVDITTSDRDKSKYGTLSRL
AVEAGFDWVYYESKAHIHCSVKA
>Trichuris suis, XKFD51835.1, 95-199
NIVFKDEEGTGADRIMTNRCRYKLNLLALLVSNFWPGVKLRVIDAWEERNRQVVGSLHYEGRAVDITTSDRDNRKIPRLARL
AVQAGFDWVYFESRQHVHASVKS
>Capitella teleta, XAAZ04357.1, 82-186
DVVFKDEEGTGADRIMSQRCKDKINTLAISVMNQWPGVKLKVTEAWDEDGFHAKDSLHYEGRAVDITTDDRDRSKYGMLARL
AVEAGFDWVYYENRGHIHCSVKS
>Pipra filicauda, XXP_027590561.1, 84-188
DIIFKDEENTGADRLMTQRCKDRLNSLAISVMNQWPGVKLRVTEGWDEDGHHSEESLHYEGRAVDITTSDRDRNKYGMLARL
AVEAGFDWVYYESKAHIHCSVKS
>Branchiostoma3 floridae, XXP_002607850.1, 269-370
HPVGFTPSQRCADRVMSKRLYTALLRVDKHVREQLNARLRITEAWDEPHSGAADGDQAENSLHYEGRAAKLELSGSSDLTSL
AKYCICADIDYVEHKGTYLF
>Acropora millepora, XXP_029199742.1, 68-170
EIDFANDDCRRMTARAKSKIDVLASRVRGRWSNVRLRVILGWTDQIPVDTQKLLHYEGRALRLQTSDRDSSKLRTLAGLAVE
AGFDWVYYASSSYIHASVIRD
>Euperipatoides kanangrensis, XVDH80594.1, 88-190
IIFKDEEGTGADRLMTQRCKEKLNTLAISVMNQWPGIKLRVTEAWDEDNHHSAESLHYEGRAVDITTSDRDRSKYGMLARLA
VEAGFDWVFYESRAHIHCSVK
~~~

